# RNA-seq library preparation from single pancreatic acinar cells

**DOI:** 10.1101/085696

**Authors:** Damian Wollny, Sheng Zhao, Ana Martin-Villalba

**Affiliations:** Molecular Neurobiology, German Cancer Research Center (DKFZ), Heidelberg, Germany

## Abstract

Single cell RNA sequencing technology has emerged as a promising tool to uncover previously neglected cellular heterogeneity. Multiple methods and protocols have been developed to apply single cell sequencing to different cell types from various organs. However, library preparation for RNA sequencing remains challenging for cell types with high RNAse content due to rapid degradation of endogenous RNA molecules upon cell lysis. To this end, we developed a protocol based on the SMART-seq2 technology for single cell RNA sequencing of pancreatic acinar cells, the cell type with one of the highest ribonuclease concentration measured to date. This protocol reliably produces high quality libraries from single acinar cells reaching a total of 5x10^6^ reads / cell and ∼ 80% transcript mapping rate with no detectable 3´end bias. Thus, our protocol makes single cell transcriptomics accessible to cell type with very high RNAse content.

## Introduction

A large number of protocols for RNA sequencing (RNA-seq) of single cells have recently been established (Picelli, 2016). Yet, library preparation from cell types, which inherently entail large amounts of RNAses can significantly impair library preparation because of RNA degradation. Thus, protocols are needed to enable transcriptomic analysis for cell types with high RNAse content, such as pancreatic acinar cells (Barnard, 1969).

The pancreas mostly consists of acinar cells that make up the vast majority of all cell types in the adult organ. Further, this cell type represents the cell-of-origin for pancreatic ductal adenocarcinoma (PDAC), one of the most malignant tumours among all types of cancer (Hidalgo, 2010). Despite the importance of studying this cell type, RNA isolation - even from large amounts - is classically considered to be very challenging (Chirgwin et al., 1979). This technical challenge has potentially precluded in-depth analysis of adult acinar cells up to this point.

Thus, we aimed at developing a protocol for whole transcriptome analysis by RNA-seq from small amounts of input RNA derived from pancreatic acinar cells. By combining chaotropic salt denaturation with solid-phase reversible immobilization technology, we enable high quality library preparation from single acinar cells. Hence, our protocol offers a low cost solution for overcoming protein mediated RNA degradation, which so far impeded high quality single cell RNA studies from RNAse rich tissues such as the pancreas, lung or spleen.

## Protocol development

For single cell RNA sequencing of acinar cells, we decided to adapt the SMART-seq2 technology based on the robust nature of the protocol according to our experience (Llorens-Bobadilla et al., 2015). We assessed the performance of library preparation conditions by microcapillary electrophoresis of preamplified cDNA. Library preparation from single acinar cells using standard conditions as previously described (Picelli et al., 2014) led to cDNA libraries of very small size and low concentration (Fig. 1a-c – “normal conditions”). The shift towards smaller fragments indicated degradation of the input material. We hypothesized that acinar derived RNAses would rapidly degrade mRNA prior to reverse transcription. Although the lysis buffer contains RNAse inhibitors, the extensive amount of RNAses will likely exceed the inhibitor concentration. In order to hold RNA degradation upon lysis, we transferred the reaction tube to liquid nitrogen directly after the cell lysis until we proceeded with cDNA synthesis. Although this increased the average cDNA length and yield, the broad peak distribution suggested persistent RNA degradation impeding full-length cDNA synthesis (Fig. 1a-c – “N2”). The secondary structure of RNAses strongly depends on disulfide bridges (Klink et al., 2000). We therefore supplemented our lysis buffer with either dithiothreitol (DTT) or β-mercaptoethanol (2ME) to reduce the disulfide bonds. The addition of either reducing agent resulted in slightly higher average library size and consistently higher cDNA yield as compared to previous buffer compositions (Fig. 1a-c – “DTT” / “2ME”). However, the peak of the library distribution was at ∼150bp indicating substantial RNA degradation (Fig. 1a). Next, we tried to target the pH dependent catalytic activity of ribonucleases. The enzyme activity of RNAse A has a pH optimum at ∼7.4, which is declining at alkaline and acidic conditions (Findlay et al., 1962). In contrast, the cDNA synthesis using M-MLV reverse transcriptase is performed at a pH of 8.3 (Kotewicz, 1988). Since the protocol does not involve purification steps between cell lysis and reverse transcription, we hypothesized that shifting the pH to 8.3 at the point of cell lysis would substantially hamper RNAse activity, but not affect reverse transcription. In contrast to all previous conditions the cDNA libraries obtained through pH shift of the lysis buffer demonstrate a peak at long fragments for a number of libraries (Fig. 1a – “pH8.3”). Although this indicates successful library preparation, the cDNA length and yield varied considerably from sample to sample (Fig. 1b,c). Thus, pH optimization might be a more attractive, low-cost approach for cDNA synthesis from tissues other than the pancreas. Previously, RNA isolation from large amounts of pancreatic tissues has been successful by making use of chaotropic agents (Chirgwin et al., 1979). In this approach, RNA degradation was prevented by lysing the cells using high concentration of guanidine hydrochloride (Chirgwin et al., 1979). Chaotropic agents such as guanidine hydrochloride or urea lead to denaturation of proteins. Subsequently, RNA could be isolated by phenol-chlorophorm extraction (Chomczynski and Sacchi, 1987). Although promising, this approach was not directly applicable to our low input protocol because an essential component of the protocol was the avoidance of purification steps before cDNA amplification (Picelli et al., 2014). A lysis buffer containing high concentrations of guanidine hydrochloride would also denature all enzymes subsequently added for reverse transcription and amplification. To circumvent this problem, we combined the chaotropic protein denaturation with solid-phase reversible immobilization (SPRI) technology. We prepared a buffer containing 6M guanidine hydrochloride and 0.2% Triton X-100 to lyse the cells and denature the released proteins. We added the reducing agent 2-mercaptoethanol to the lysis buffer as a result of the small but consistent enhancement in cDNA yield and length that we observed (Fig. 1a-c – “GuaHCl + 2ME”). After cell lysis we directly transferred the reaction tube into liquid nitrogen for the time of cell collection (Fig. 2). Next, we added SPRI beads to the lysate. SPRI beads are carboxyl-coated magnetic particles, which are dissolved in polyethylene glycol (PEG) (Fig. 2) (Hawkins et al., 1994). Application of high PEG concentrations leads to the precipitation of RNA and DNA onto the carboxylated beads at nearly 100% efficiency (Hawkins et al., 1994). Subsequent washing steps remove the guanidine hydrochloride containing lysis buffer as well as acinar cell derived RNAses (Fig. 2). It should be noted that PEG could also be used for protein precipitation (Lis, 1980). However, efficient protein precipitation onto carboxylated beads requires acetonitrile treatment (Hughes et al., 2014). Therefore, residual RNAses, which are inadvertently carried over by PEG precipitation, might be not sufficient for degradation of RNA. After the final washing steps, the elution step was avoided to trap RNA on the beads for cDNA synthesis as previously described (Fig. 2) (Shalek et al., 2013). The protocol led to highly increased average cDNA length and yield with very limited sample-to-sample variablilty (Fig. 1a-c “GuaHCl + 2ME”). We observed indistinguishable libraries by using dithiothreitol instead of β-mercaptoethanol,which strongly indicated successful and robust cDNA library preparation from single acinar cells. Hence, we continued the library preparation from samples lysed using guanidine hydrochloride with and without reducing agents (GuaHCl ± DTT / 2ME) and sequenced them on an Illumina HiSeq2000 sequencer (paired-end 100bp) as previously described (Llorens-Bobadilla et al., 2015).

**Figure 1.**
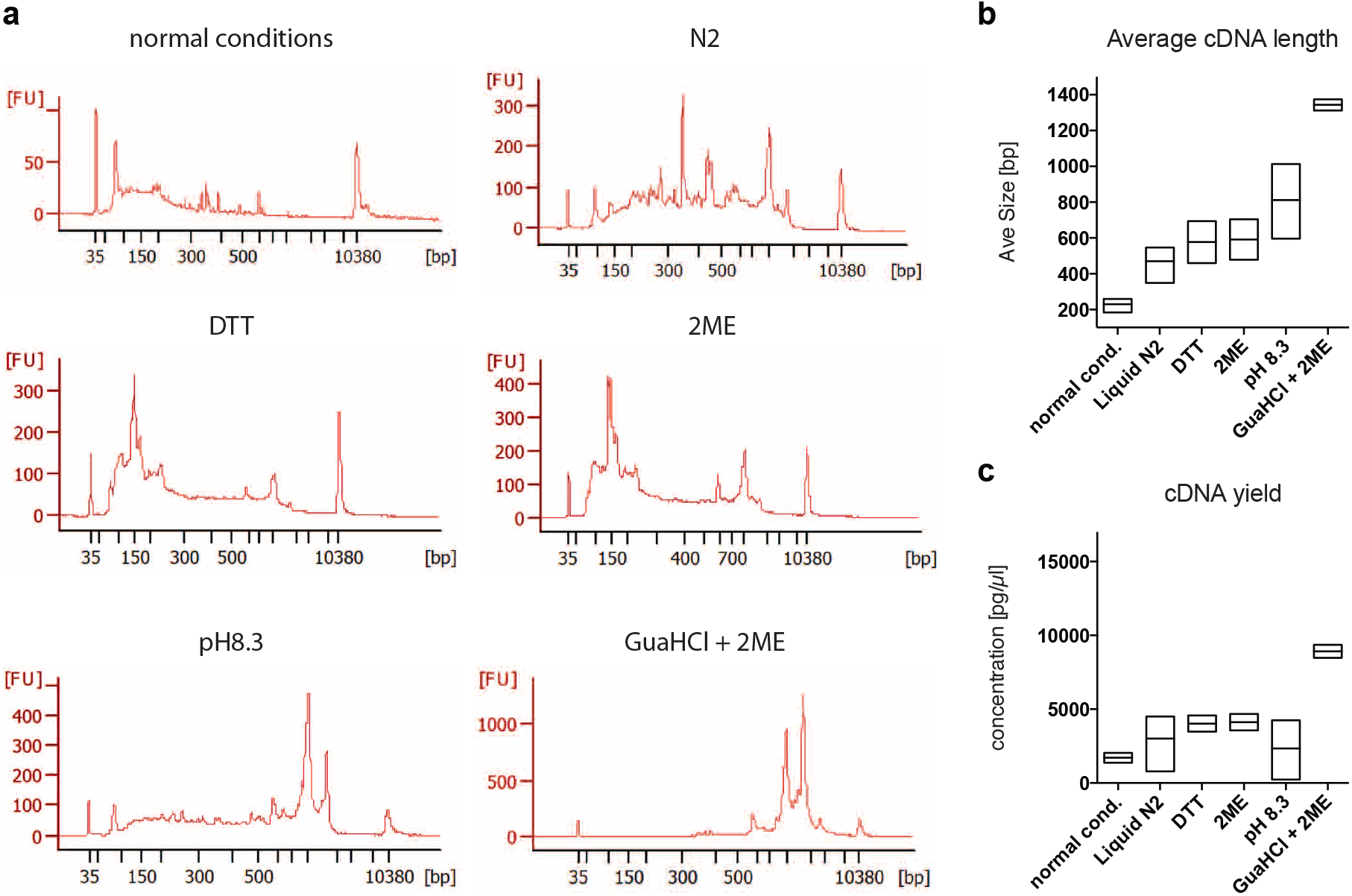
Protocol development for cDNA library preparation from single acinar cells. (**a**) Representative Bioanalyzer profiles of cDNA libraries obtained from single acinar cells under varying cell lysis conditions. (**b**) Average size of cDNA library length obtained under conditions described in (a). (**c**) cDNA yield produced under conditions described in (a).

**Figure 2.**
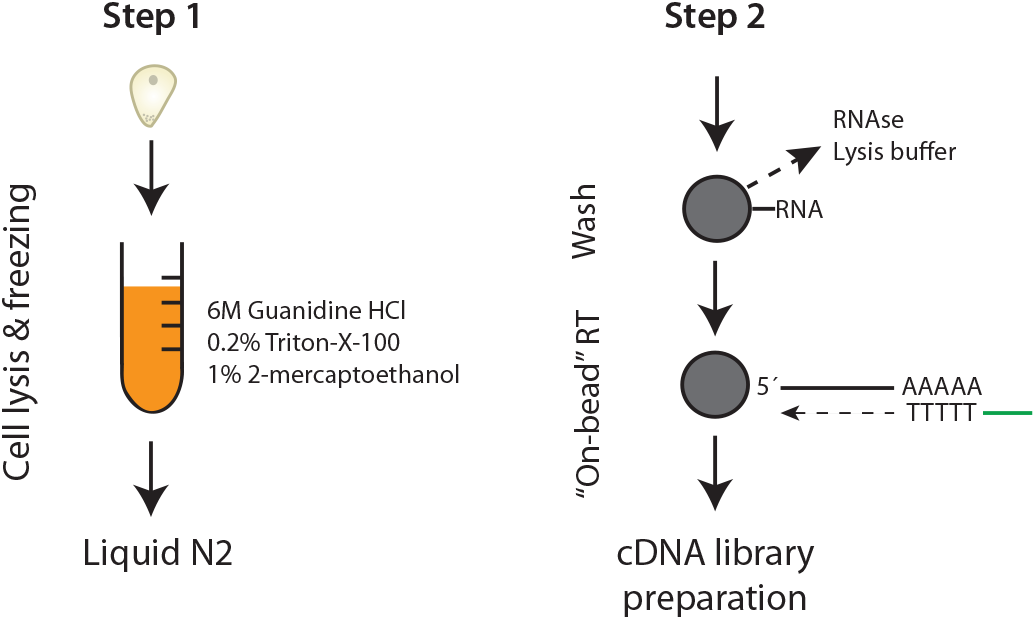
cDNA library preparation protocol for single acinar cells. Schematic representation of the step-by-step protocol for cDNA library production from single acinar cells. Grey circles represent SPRI beads.

Bioinformatic quality control analysis of the sequenced libraries revealed an overall high mapping rate and a high total amount of reads comparable to previously published single cell RNA sequencing studies (Table 1) (Llorens-Bobadilla et al., 2015; Treutlein et al., 2014). Further, we observed good overall coverage along the transcripts without 3´ bias (Fig. 3).

**Figure 3.**
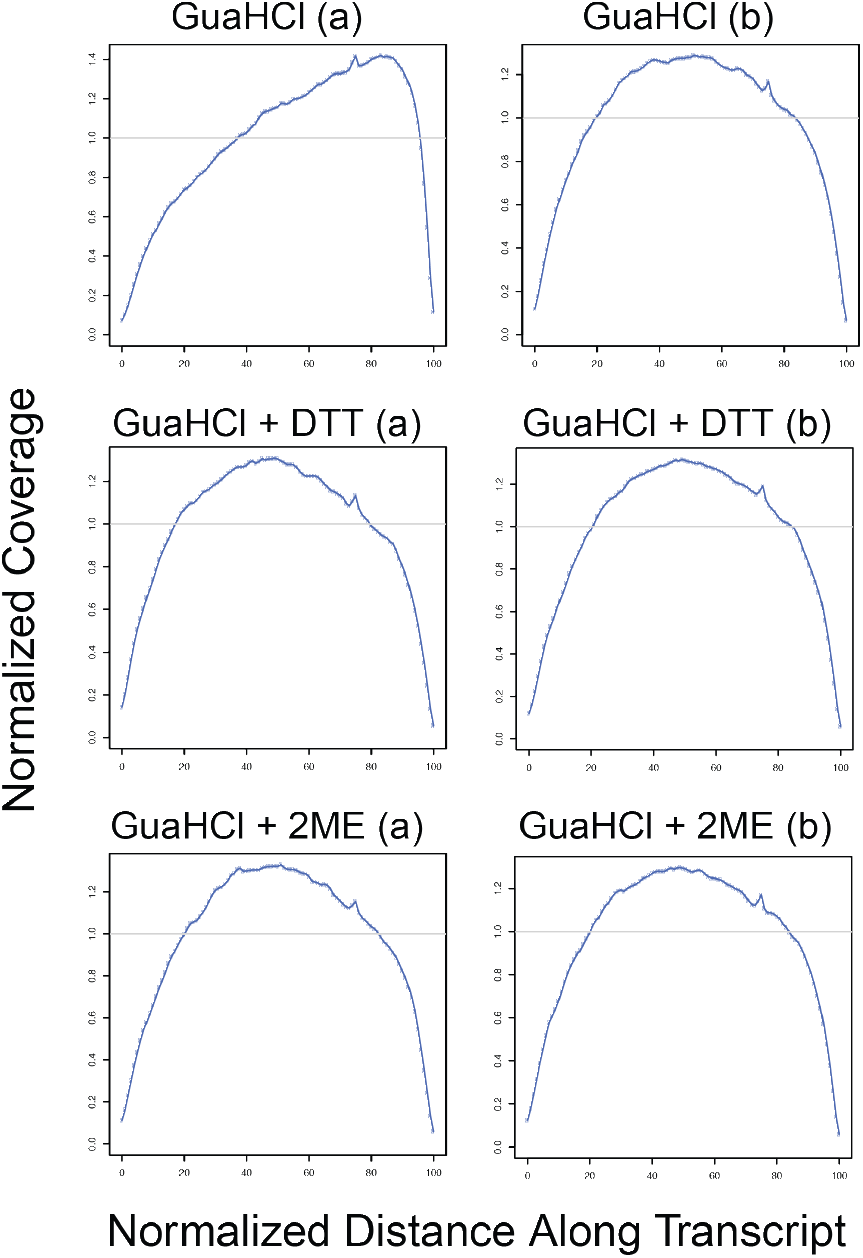
Normalized 3 ́ and 5 ́ coverage of sequenced reads along the transcripts. Analysis of 3´-5´coverage of libraries prepared from single acinar cells using the described lysis buffer conditions. (a) and (b) represent library replicates from two independent acinar cells.

**Table 1.** Quality control of single acinar cell libraries

| Lysis Buffer | Total reads / cell | Mapping rate [%] |
| --- | --- | --- |
| GuaHCl (a) | 5,779,349 | 64.56 |
| GuaHCl (b) | 4,693,346 | 81.50 |
| GuaHCl + DTT (a) | 5,011,597 | 81.28 |
| GuaHCl + DTT (b) | 4,946,490 | 81.93 |
| GuaHCl + 2ME (a) | 5,414,012 | 85.36 |
| GuaHCl + 2ME (b) | 5,627,489 | 83.57 |

Blue highlight denotes library with low mapping rate and 3´ transcript coverage bias (Fig 3). (a) and (b) represent library replicates from two independent acinar cells. GuaHCl % guanidine hydrochloride; DTT % dithiothreitol; 2ME % β-mercaptoethanol

Among the tested lysis buffer conditions, one library prepared without reducing reagents showed markedly decreased mapping rate (Table 1 – GuaHCl (a)) as well as a 3´ transcript coverage bias (Fig. 3 – GuaHCl (a)). This result indicates that, despite the high concentration of guanidine hydrochloride, reducing agents will benefit successful library preparation. Between the two reducing agents used, addition of β-mercaptoethanol resulted in slightly higher yield of total reads per cell and mapping rate compared to dithiothreitol (Table 1). Hence, we decided to supplement our lysis buffer using β-mercaptoethanol. Furthermore, we tested expression levels of digestive enzymes, which are expressed at high levels in acinar cells (Uhlen et al., 2015). We find high expression of these genes across all analysed acinar cells, confirming the acinar identify of the tested cells (Fig. 4). However, in agreement with the quality control analysis, we observed consistently decreased enzyme expression in one of the samples prepared without reducing conditions during cell lysis (Fig. 4 – GuaHCl (a)). Hence, this analysis again confirms the importance of supplementing the lysis buffer with reducing reagents.

**Figure 4.**
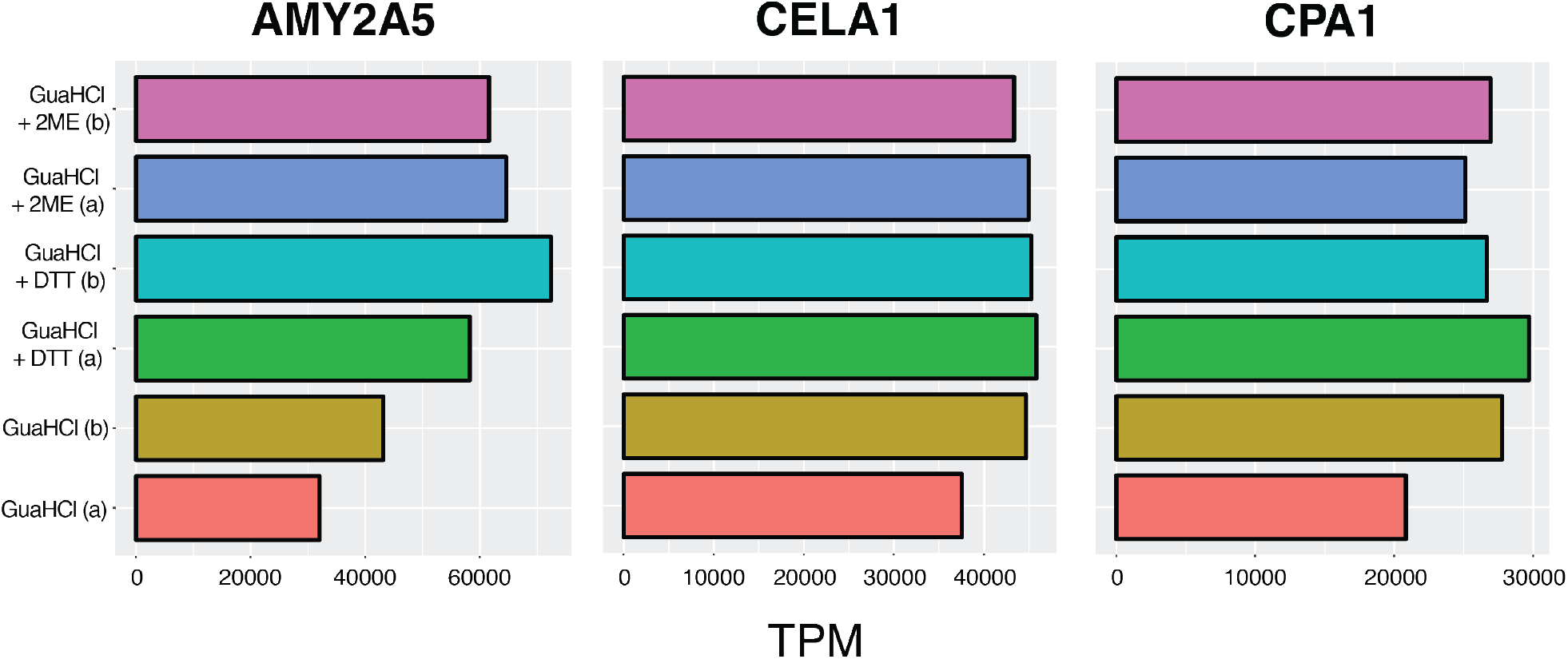
Expression of acinar cell markers. Expression values of digestive enzymes amylase (*Amy2a5*), elastase (*Cela1*), carboxypeptidase A1 (*Cpa1*) highly expressed in pancreatic acinar cells. TPM, transcripts per million.

The final protocol as described in Fig. 2 enabled successful library preparation from over 100 single acinar cells (Wollny et al., 2016). Thus, our protocol provides the opportunity for single cell sequencing from cell types with high endogenous expression of ribonucleases.

## Materials

### Chemicals & Reagents

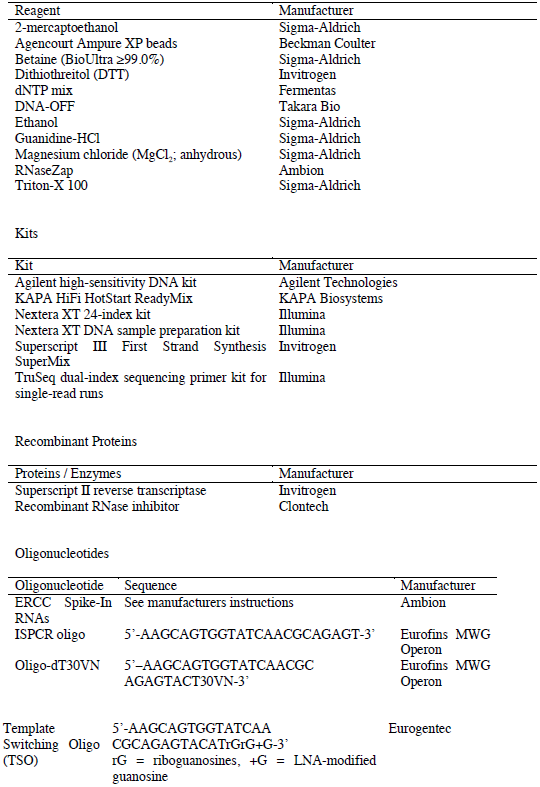

## Protocol

The protocol described below is based on the SMART-seq2 library preparation protocol (Picelli et al., 2014). We therefore recommend getting familiar with the SMART-seq2 protocol before implementing our adjustments to this protocol. Standard precaution for RNA work (disposable gloves, clean work surface, dust mask etc.) should be adhered to during all steps of the protocol.

1. Prepare lysis buffer containing 6M Guanidine HCl, 0.2% (vol/vol) Triton X-100 and 1% (vol/vol) 2-mercaptoethanol. Transfer 4µl of lysis buffer mix in every tube for single cell lysis.
2. Isolate pancreatic acinar cells as previously described (Wollny et al., 2016).
3. Pick single acinar cell (resuspended in DPBS without Mg2+ and Ca2+) in a volume of 0.5µl and directly transfer in 4µl lysis buffer. Put the tube in liquid N2 immediately. Note: Acinar cells are very adhesive and tend to re-aggregate. Handpicking using a microscope is strongly recommended in order to avoid collection of doublets or cell aggregates. Further, we tried to limit the time of cell collection to a maximum of 60 min in order to minimize dissociation induced transcriptional changes.
4. After collecting all cells, add 2.2x SPRI beads (Agencourt RNAclean XP from Beckman Coulter – need to be at room temperature). Incubate for 5 min at room temperature. Wash beads 2x using 70% EtOH as described in the manufacturer´s protocol. Completely remove EtOH traces and let the beads dry.
5. Resuspend dried beads using a solution containing 2μl of nuclease-free H2O, 1μl of Oligo-dT30VN primer and 1μl of dNTP mix. Use Oligo-dT30VN and dNTP concentration and sequence as described in (Picelli et al., 2014). Incubate beads including oligo-dT and dNTP for 3min at 72°C. Optional: ERCC Spike-In RNA mix (1:10,000 - Ambion) may be added to the Oligo-dT30VN / dNTP mix.
6. Add reagents for reverse transcription agents (5.5µl) to the beads and run reverse transcription program, both as described in (Picelli et al., 2014) – step 9-11.
7. Leave the beads inside the tube and add PCR mix and run PCR cycles as in (Picelli et al., 2014) (18 cycles).
8. Clean up PCR product by adding 0.8x DNA SPRI beads (Agencourt Ampure XP) to the PCR mix. Resuspend dried beads in 15µl clean H2O and run 1µl on a high sensitivity DNA Bioanalyzer chip. Note: We tested every cDNA library on a Bioanalyzer chip by manually examination of the bioanalyzer profile before proceeding with tagmentation. Depending on the experience and speed of the experimenter we obtained a success rate for library preparation of ˜ 60-80% (data not shown).
9. Continue library preparation according to the SMART-seq2 protocol as previously described (Picelli et al., 2014; Llorens-Bobadilla et al., 2015)

## Acknowledgement

We would like to thank Simone Picelli and Alex Shalek for helpful advice during establishment of this protocol. Further, we would like to thank the members of the Martin-Villalba lab for support and useful suggestions. We thank the NCT Heidelberg, the Helmholtz alliance preclinical comprehensive cancer center, the DFG (SFB873), and the DKFZ for funding.

